# Distinct VIP interneurons in the cingulate cortex encode anxiogenic and social stimuli

**DOI:** 10.1101/2020.12.22.424056

**Authors:** Connor Johnson, Lisa N. Kretsge, William W. Yen, Balaji Sriram, Jessica C. Jimenez, Tushare J. Jinadasa, Alexandra O’Connor, Ruichen Sky Liu, Thanh P. H. Nguyen, Eun Seon Cho, Erelle Fuchs, Eli D. Spevack, Berta Escude Velasco, Frances S. Hausmann, Alberto Cruz-Martín

**Author notes:** Co-first author (Connor Johnson, Lisa N. Kretsge, William W. Yen). Corresponding author Corresponding author contact information: Name: Alberto Cruz-Martín, Postal address: Boston University, 24 Cummington Mall, Room 426, Boston, MA 02215.

## Abstract

A hallmark of higher-order cortical regions is their functional heterogeneity, but it is not well understood how these areas encode such diverse information. The anterior cingulate cortex (ACC), for example, is important in both emotional regulation and social cognition. Previous work shows activation of the ACC to anxiety-related and social stimuli, but it is unknown how subpopulations or microcircuits within the ACC simultaneously encode these distinct stimuli. One type of inhibitory interneuron, which is positive for vasoactive intestinal peptide (VIP), is known to alter the activity of many cells in local cortical microcircuits, but it is unknown whether the activity of VIP cells in the ACC (VIP^ACC^) encodes anxiety-related or social information. Using in vivo calcium imaging and miniscopes in freely behaving mice to monitor VIP^ACC^ activity, we identified distinct, non-overlapping subpopulations of VIP^ACC^ that preferentially activated to either anxiogenic, anxiolytic, social, or non-social stimuli. We determined that stimulus-selective cells encode the animal’s behavioral states and VIP interneuron clusters may co-activate, improving this encoding. Finally, we used trans-synaptic tracing to show that VIP^ACC^ receive widespread inputs from regions implicated in emotional regulation and social cognition. These findings demonstrate not only that the ACC is not homogeneous in its function, but also that there is marked functional heterogeneity even within disinhibitory interneuron populations. This work contributes to our understanding of how the cortex encodes information across diverse contexts and provides insight into the complexity of neural processes involved in anxiety and social behavior.

## INTRODUCTION

Cortical subregions are often implicated in a variety of behavioral functions, but it is not well understood how these areas encode such diverse information. The anterior cingulate cortex (ACC) is necessary for emotional processing and social cognition, but how it encodes stimuli relevant to both processes is unknown (1–6). In humans, ACC activity increases when healthy subjects perform social tasks and is higher in anxiety disorder patients who demonstrate increased symptom severity (7). In rodents and non-human primates, ACC inhibition via chemical lesions or genetic manipulations impairs social behaviors (8–10). In addition, stimulating activity or silencing a cytoskeletal protein in the ACC alters anxiety-like behaviors (11, 12). Recent studies in rodents have provided insight into the ACC’s importance in emotional and social behaviors by monitoring either single unit or bulk activity of this region (6, 8, 11, 13–16). However, these data do not parse out the roles of different neural subtypes or microcircuits within the ACC. It remains unknown how neuronal representations of diverse stimuli are embedded within ACC subcircuits.

Inhibitory neurons that are positive for vasoactive intestinal peptide (VIP) inhibit other inhibitory cells, thereby driving excitatory pyramidal cell (Pyr) activity (17–20). VIP interneurons, therefore, are in a unique position to modulate local activity (21, 22). They also receive long-range inputs from other brain regions, which may allow them to coordinate the activity of the ACC with other brain regions to respond to diverse stimuli (18, 20, 23). In the hippocampus, VIP interneurons form functional clusters that are differently modulated by behavioral states, but this has not been studied in higher-order cortical regions like the ACC (24). Acetylcholine and serotonin can modulate functionally distinct groups of VIP interneurons and may differentially engage them during diverse behaviors (25–27). Previous work shows that cortical VIP cells exhibit diverse molecular, morphological, and electrophysiological properties. However, it is unknown whether VIP cells have heterogeneous functions in the ACC and whether disinhibitory circuits are involved in ACC processing of diverse stimuli.

We used miniscopes to perform in vivo single-cell resolution calcium imaging of VIP interneurons in the ACC (VIP^ACC^) to investigate their functional heterogeneity while mice performed tasks to assay anxiety-like behaviors, general sociability and social novelty. We identified distinct VIP^ACC^ subgroups that reliably encoded behavioral states by preferentially activating to anxiety-related, social, or non-social stimuli. In addition, we found that averaging selective cell activity made this coding more reliable. These results suggest that the improvement in coding was mediated through co-activation of these inhibitory subpopulations. When the same neurons were monitored across anxiety-related and social tasks, we determined that the majority of VIP^ACC^ that were engaged during these tasks were highly selective, activating to only one specific stimulus. Lastly, using rabies trans-synaptic mapping, we showed that VIP^ACC^ receive inputs from brain regions implicated in emotional regulation and social behavior. Our data show that VIP^ACC^ are functionally heterogeneous and that non-overlapping subgroups of VIP^ACC^ activate preferentially, providing a cellular substrate for encoding different types of stimuli.

## MATERIALS AND METHODS

### Animals

Animals were grouped housed in a 12-hr light/dark schedule vivarium with food and water ad libitum. Experimental mice were male postnatal day 60-120 VIP-Cre mice (Vip-IRES-cre, #010908, The Jackson Laboratory, Bar Harbor, Maine (28)). Stimulus mice for social interaction were littermates (male VIP-Cre) or novel (age matched male CD-1 IGS, strain code: 022, Charles River Laboratories, Wilmington, Massachusetts). All procedures were approved by the Institutional Animal Care and Use Committee (IACUC) at Boston University and practices were consistent with the Guide for the Care and Use of Laboratory Animals and the Animal Welfare Act.

### Surgeries

Surgeries were performed using aseptic surgical techniques with autoclaved instruments. Animals were weighed and anesthesia was induced in a chamber with an isoflurane–oxygen mixture (4% [v/v]). Anesthesia was maintained throughout the procedure via mask inhalation of an isoflurane– oxygen mixture (Henry Schein, Melville, New York, 1-1.5% [v/v]). Animals were kept on a heating pad (T Pump, Gaymar Industries Inc., Orchard Park, New York) for the duration of the surgeries and for 30-min recovery periods before being returned to their home cages. Animals were injected with buprenorphine (3.25 mg/kg; SC, Patterson Veterinary, Greeley, Colorado), meloxicam (5 mg/kg; SC, Covetrus, Dublin, Ohio), and dexamethasone (Henry Schein, 2.5 mg/kg; SC) and the fur on the top of the head was removed. Animals were head-fixed using a stereotax (Kopf Instruments, Tujunga, California). The surgical area was sterilized with 10% povidone-iodine and 70% isopropyl alcohol (CVS, Woonsocket, Rhode Island) and local anesthetic was applied (lidocaine 1% and epinephrine 1:100,000; SC, Henry Schein). After surgeries, post-operative analgesics were administered for 2 days, twice per day (buprenorphine 0.01 mg/kg; SC and meloxicam 5 mg/kg; SC).

#### Viral injections

After the preparations above, an incision was made in the skin along the midline of the skull. A craniotomy was made over the injection site using a pneumatic dental drill (eBay, Inc., San Jose, California). Using a stereotax and the Nanoject II (Drummond Scientific, Broomall, Pennsylvania), a pulled-glass pipette (BF150-117-10; tip size ~3-15 μm, Sutter Instrument Co., Novato, California) was lowered into the ACC (AP: +0.90 mm, ML: −0.30 mm, DV: −1.00 mm) and virus was injected. After pipette removal, the skin was sutured with non-absorbable sutures (AD Surgical, Sunnyvale, California).

To monitor VIP^ACC^ activity, we injected 460 nl of an adeno-associated virus (AAV) that expresses GCaMP6f, a fluorescent Ca^2+^ indicator, in a Cre-dependent manner (AAV9-CAG-Flex-GCaMP6f.SV40, titer: 5.23 × 10^13^ GC/ml, packaged by the University of Pennsylvania Vector Core, Philadelphia, Pennsylvania).

For trans-synaptic tracing of monosynaptic inputs to VIP^ACC^, we first injected 128 nl of a helper AAV that expresses target proteins under the human synapsin-1 promoter: AAV2/1-synP-Flex-splitTVA-EGFP-B19G (AAV-TVA-Glyco) (29) (titer: 0.98 × 10^12^ GC/ml, University of North Carolina Viral Core, Chapel Hill, North Carolina). This AAV contained genes to express EGFP, the avian sarcoma/leukosis virus subtype A receptor (TVA, which confers infection capability to rabies virus pseudotyped with the avian sarcoma leucosis virus glycoprotein (EnvA)), and the rabies virus glycoprotein (G) (29, 30) (which is necessary for trans-synaptic transport of glycoprotein gene-deleted (ΔG) rabies virus (RV)) (23). These three genes were in frame and separated by porcine teschovirus self-cleaving 2A elements (29). After allowing the virus to express for one month (31), the skin was re-incised, a new craniotomy was drilled, and 128 nl of ΔG EnvA pseudotyped RV with mCherry was injected (RV*dG*, titer: 1.5 × 10^9^ GC/ml, Boston’s Children Hospital Viral Core, Boston, Massachusetts). There is no cognate receptor for EnvA in the mouse, so RV*dG* only infects TVA-expressing cells. Together, Glyco, TVA, and RV*dG* allow for retrograde monosynaptic tracing only from Cre-expressing cells. In Cre-negative mice (N = 6), we performed these injections and saw no labeled cells, suggesting a lack of leaky expression (data not shown).

#### GRIN lens implants

To image neuronal activity with a miniaturized microscope (miniscope), a gradient-index (GRIN) lens was implanted in the ACC at least 2 weeks after viral injection (to allow for maximal GCaMP6f expression). After the preparations above, the scalp was re-incised and a 1 mm diameter craniotomy was drilled, centered around the viral injection. Three screws (Fine Science Tools Inc., North Vancouver, Canada) were inserted into the skull and a layer of super glue (cyanoacrylate, Krazy glue, High Point, North Carolina) was applied to ensure the lens and dental cement adhered strongly. Dura over the ACC and a small region of the secondary motor cortex were aspirated using a blunted 18G needle (BD, Franklin Lakes, NJ) coupled to a vacuum line. A GRIN lens (Table S1) attached to a stereotax via custom 3D-printed implant assembly (Fig S1) was lowered into the ACC at a 20° angle (AP: +0.90 mm, ML: −0.12 mm, DV: −0.13 mm), to improve access to the ACC and minimize the risk of puncturing the midline vasculature. Lenses were adhered to the skull with optical glue (Norland Products Inc., Cranbury, NJ) and dental cement (Ortho-Jet™ Liquid, Black, Lang Dental, Wheeling, Illinois). An antibiotic was administered via the water supply (Biomox, 0.75mgl/ml, Henry Schein) for 10 days after surgery.

### 3D-printed miniscopes and baseplating

3 weeks after lens implant surgeries (to allow for recovery and optimal GCaMP6f expression), animals were anaesthetized, as described above, and GCaMP6f expression was assessed by imaging the fluorescent signal using miniscopes. When GCaMP6f-positive neurons were visible, baseplates were attached to the skulls with dental cement using a custom 3D printed baseplating assembly (Fig S1E-F). The miniscopes used were modified from two existing designs (Ghosh et al. 2011 (32), Liberti et al. 2017 (33)) to allow it to detach from the baseplate (Fig S1), which made it possible to co-housed the animals without risking miniscope damage. Miniscopes were attached to baseplates at a 15-20 degree angle relative to the midline to align with the GRIN lenses (Fig S1B). Miniscope models are available at https://github.com/CruzMartinLab. Commercially available parts are listed in Table S1. Custom parts were 3D printed (Form 3 Printer, Black Resin FLGPBK03, Formlabs, Somerville, Massachusetts) and assembled in-house (Fig S1).

### Perfusions and Histology

Animals were perfused to visualize viral injections and lens placements. Mice were injected with an overdose of sodium pentobarbital (250 mg/kg; IP, Vortech Pharmaceuticals Ltd., Dearborn, Michigan) and transcardially perfused with phosphate buffered saline (PBS) followed by 4% paraformaldehyde in PBS (PFA). Brains were extracted, stored in PFA for 24 hr at 4°C, and then transferred to a 30% (w/v) sucrose solution for 48 hr at 4°C. Tissue was sectioned at 50-100 μm using a freezing sliding microtome (SM2000, Leica Biosystems, Buffalo Grove, Illinois). Sections were mounted onto slides (Globe Scientific Inc., Mahwah, New Jersey) using Fluoromount-G mounting medium with DAPI (Thermo Fisher Scientific, Waltham, Massachusetts) to visualize nuclei.

### Imaging

Sections were imaged using an upright wide-field microscope (Nikon Eclipse Ni, Nikon Instruments Inc. Melville, New York) controlled by NisElements (Nikon Instruments Inc., 4.20). Images were acquired using a Plan Fluor 4X (NA 0.13) or 10X (NA 0.3) objective with standard Nikon HQ filter cubes for DAPI, EGFP/GCaMP, and mCherry. Images were analyzed as TIFFs in ImageJ (National Institute of Health, Bethesda, Maryland) and compared to the Allen Mouse Brain Atlas (34) (brain-map.org/api/index.html) to identify brain regions.

### Behavioral assays and in vivo Ca^2+^ imaging

Before any behavioral testing, mice were handled for 10 min for 3 days to acclimate them to the experimenter. For anxiety-related assays, mice were not exposed to the arenas prior to testing, but for the social task, implanted mice were acclimated to the arena with empty cups for 10 min per day for 2 days prior to the task. This ensured that, during the social behavioral tasks, mice were familiar with the empty cups. In addition, stimulus mice were acclimated to being housed in cups for 10 min on each of these days. Arenas were custom made from acrylic and HDPE (McMaster-Carr, Elmhurst, Illinois) and were cleaned with 70% ethanol between trials and animals. Behavior was recorded (C270 Webcam, Logitech, Lausanne, Switzerland) at 30 frames/s under overhead lighting (20 lux). After miniscopes were attached, animals were given 10 min to rest in an empty chamber before we started the experiments. Ca^2+^ videos were acquired at 20 frames/s using a Miniscope Data Acquisition PCB and Data Acquisition Software ((35)Table S1). We acquired images with a field of view of 720 × 480 pixels (approximately 800 μm × 600 μm). Excitation LED power was adjusted to optimize imaging for each animal with a maximum output of 1 mW. To avoid bleaching GCaMP6f, no trials were longer than 10 min.

#### Elevated zero maze (EZM)

To assay anxiety-like behavior, we used the EZM, an elevated circular arena with two open arms and two closed arms (track diameter = 50 cm, track width = 5 cm, wall height for closed arms = 40 cm, height of track = 61 cm). Open arms are considered anxiogenic and closed are anxiolytic. Mice were initially placed in the closed arm and recorded for 10 min (36).

#### Social task

The arena contained two mesh wire cups – one on each end of the arena (50 × 25 × 30.5 cm length-width-height). On Day 1 (Sociability), one mesh wire cup housed a littermate and the other was empty. On Days 2 and 3 (Social Novelty), one cup housed a littermate and the other housed an entirely novel male CD-1. Each day the experimental mouse was placed in the center of the arena and given 10 min to explore the arena and cups. The side of the arena with each stimulus was randomized.

#### Open field (OF)

To assay anxiety-like behavior and locomotion, the OF was used. The center of the arena is considered anxiogenic and the periphery is anxiolytic. Mice were placed in the center of a custom-made acrylic arena (50 × 50 × 30 cm length-width-height) and allowed to explore for 10 min.

### Behavioral analysis

Behavior was analyzed using DeepLabCut (37), an open-source program that uses machine learning to track body parts in behavioral videos, as described by Comer et al. (38). To assess accuracy, videos labeled by the software were inspected by a trained observer and custom MATLAB (MathWorks, Natick, Massachusetts) scripts were used to verify that DeepLabCut located each body part at least 95% of the total time the animal was tracked. For EZM and OF, we tracked the centroid of the mouse’s body to determine velocity, distance traveled, and when the mouse was in each zone of the arena. For the OF, we divided the arena into 25 squares (10 cm × 10 cm each) and defined the periphery as the outermost 16 and the remaining 9 as the center. For the social tasks, we tracked the head to determine close proximity to either cup. Binary behavior matrices (vectorized behavior) indicating the location of the animal were created from DeepLabCut using custom MATLAB scripts.

### Ca^2+^ imaging analysis

Ca^2+^ imaging data were processed using CaImAn (39) written in Python (https://www.python.org/). Ca^2+^ recordings underwent piecewise rigid motion correction using patches of 48 × 48 pixels with 24 × 24 pixel overlap. Following image stabilization, cell detection and extraction of df/f traces were performed with a merging threshold of activity correlation greater than 0.7 between nearby cells, and a 2.5 minimum threshold for the signal to noise ratio. df/f traces and spatial information were exported and saved in .mat format using SciPy (40). All subsequent analysis was performed using custom MATLAB and Python functions. Raw Ca^2+^ traces were z-scored using the mean baseline df/f and sigma from the entire time series for each trial. Z-scored traces report deltaf/f (df/f) values in units of standard deviation (SD). Due to the different acquisition frequencies of behavioral and Ca^2+^ imaging videos, data were aligned using timestamps from acquisition and custom MATLAB scripts. Gaps in neural data due to dropped frames (less than 1% of the total frames) were filled with averaged z-scored df/f values from surrounding frames. Starts and ends of behavioral epochs were matched to Ca^2+^ data timestamps to isolate neural activity during select behaviors.

#### Calcium activity

For all figures, Ca^2+^ activity refers to the area under the curve. To quantify this value, we used z-scored df/f traces and took their integral using the MATLAB trapz function. For all figures, excluding Fig S2I-J, area under the curve was calculated for 5 s intervals, which was chosen based on average transient length. When calculating the average area under the curve across velocities (Fig S2I-J), the intervals were 1 s. These integral values were calculated for each cell to get the Ca^2+^ activity values. When average Ca^2+^ activity is reported, that refers to the average area under the curve for all cells for each mouse.

#### Single cell ROC analysis

Responses of individual cells during different behavioral conditions were assessed within each behavioral trial using receiver operating characteristic (ROC) analysis, as previously described (41). The ROC curve demonstrates how well a single neuron’s activity matches an animal’s behavioral state, which can be quantified by calculating the area under the ROC curve (auROC) (41, 42). For each neuron in each behavioral condition, an ROC curve was generated using the true positive rate (TPR) and false positive rate (FPR) values for that cell and behavioral state. TPR and FPR were calculated across multiple binary thresholds applied to z-scored df/f traces of each cell, ranging from the minimum to maximum values of the Ca^2+^ signal. For each threshold, binarized df/f traces were compared to the binary behavioral vectors, which used binary values to indicate an animal’s presence or absence in a specific zone of the arena. TPR and FPR were then plotted against each other to create each ROC curve and auROC was calculated.

To classify cells as stimulus-selective or neutral, we determined whether the cell’s auROC value for a given stimulus was high enough to suggest it preferentially activated to a stimulus. To do this while accounting for any random alignment in our data, we calculated 1000 null values for each cell by applying circular permutations of randomized lengths to the Ca^2+^ data and calculating auROC for each of these randomized versions of the data. Cells were considered selective for a certain stimulus if the auROC of that cell was at least 2 SD greater than the mean of the null distribution (auROC > 97.5^th^ percentile). Otherwise, cells were considered neutral. For Fig 3, to calculate the auROC of “super cells”, we averaged the z-scored df/f traces of all selective cells and re-calculated auROC for that averaged data.

#### auROC analysis across tasks

For Fig 4A-D, to determine if the same cells were responsive to stimuli in different tasks, we registered cells across EZM and Sociability using CaImAn (39) (127 registered VIP^ACC^ from all 6 animals). For each cell and behavioral condition, auROC was calculated and cells were considered selective for multiple conditions (as described above) if they were selective for different stimuli across these tasks.

#### auROC analysis validation

To validate the auROC analysis (Fig S5), we looked for consistent activity changes in the cells we identified as selective. We calculated auROC values using the first half of the EZM or Sociability data, rather than the entire dataset, to identify selective cells (Fig S5A). Next, using these classifications of selectivity, we assessed Ca^2+^ activity from the second half of the data under the cell’s preferred and non-preferred conditions (Fig S5A). The preferred condition was the one that cell was selective for, whereas non-preferred was the other context or stimulus in that task.

#### Activity heatmap

For Fig S4D-E, the activity heatmap was plotted to visualize the average cell activity in 5×5 pixel spatial bins across the OF arena. Z-scored df/f traces from individual cells were normalized to their maximum value and matched with DeepLabCut centroid coordinates at the closest timestamp.

### Analysis of rabies tracing data

For retrograde mapping experiments, brains were scanned to identify signal from starter and retrogradely labeled input neurons. Cells were quantified using ImageJ. Starter cells were defined as cells that were positive for both GFP (from AAV-TVA-Glyco) and mCherry (from EnvA-ΔG-mCherry). We confirmed with DAPI that starter and retrogradely-labeled cells had a nucleus. Each animal had a different number of starter cells, so to normalize our data, we divided the number of retrogradely-labeled neurons in each region by the number of starter cells for that mouse (Inputs per starter cell). The number and location of labeled neurons in a given region was independently confirmed by 3 trained scientists. After quantifying all cells, input brain regions were divided into quartiles by number of input neurons. Only brain regions in the top two quartiles were graphed and included in Fig 4. We did not include RVdG inputs at the site of the ACC injection since leakage in TVA expression could lead to Cre-independent local labeling (43).

### Statistical analysis

Statistical analyses were performed using Graph Pad Prism 8.0 (GraphPad Software Inc., San Diego, California). For figure preparation, CorelDRAW Graphics Suite X8 (Corel Corporation, Ottawa, Canada) and ImageJ were used. The threshold for significance was set to α = 0.05 and \**p*<0.05, \*\**p*<0.01, \*\*\**p*<0.001, \*\*\*\**p*<0.0001. Data are presented as mean ± SEM, unless otherwise noted. t-tests and ANOVAs followed by appropriate post tests were used and are specified in the figure legends. For Fig S3A-B, Fig S4K-L, and Fig S6A-D, frequency distributions were fitted with Gaussians and percentages of selective cells are represented in pie charts. For all behavioral experiments, N = 6 implanted mice for Ca^2+^ imaging and N = 5 control mice that underwent no surgeries. For EZM, n = 345 cells, for Sociability, n = 310 cells, for Social Novelty Day 2, n = 350 cells, Day 3, n = 232 cells, and OF, n = 273 cells. For tracing experiments, N = 3 mice with n = 691 starter cells and 10107 retrogradely-labeled input cells.

## RESULTS

### Distinct VIP^ACC^ interneurons preferentially encode anxiogenic or anxiolytic contexts

To determine whether VIP^ACC^ are a heterogeneous population, we imaged their activity as mice performed behavioral tasks. To do this, we injected AAV9-flex-GCaMP6f and implanted GRIN lenses into the ACC of VIP-Cre mice (Fig 1A and B). Post-mortem histology showed that injections and lenses were successfully targeted to the ACC (Fig 1B), allowing monitoring of VIP^ACC^ with single-cell resolution during behavioral assays (Fig S2A, Fig 1C). 3D-printed miniscopes that detach from baseplates were adapted from two previous designs (Ghosh et al. 2011 (32), Liberti et al. 2017 (33)), assembled in-house, and used for Ca^2+^ imaging (Fig S1).

**Figure 1.**
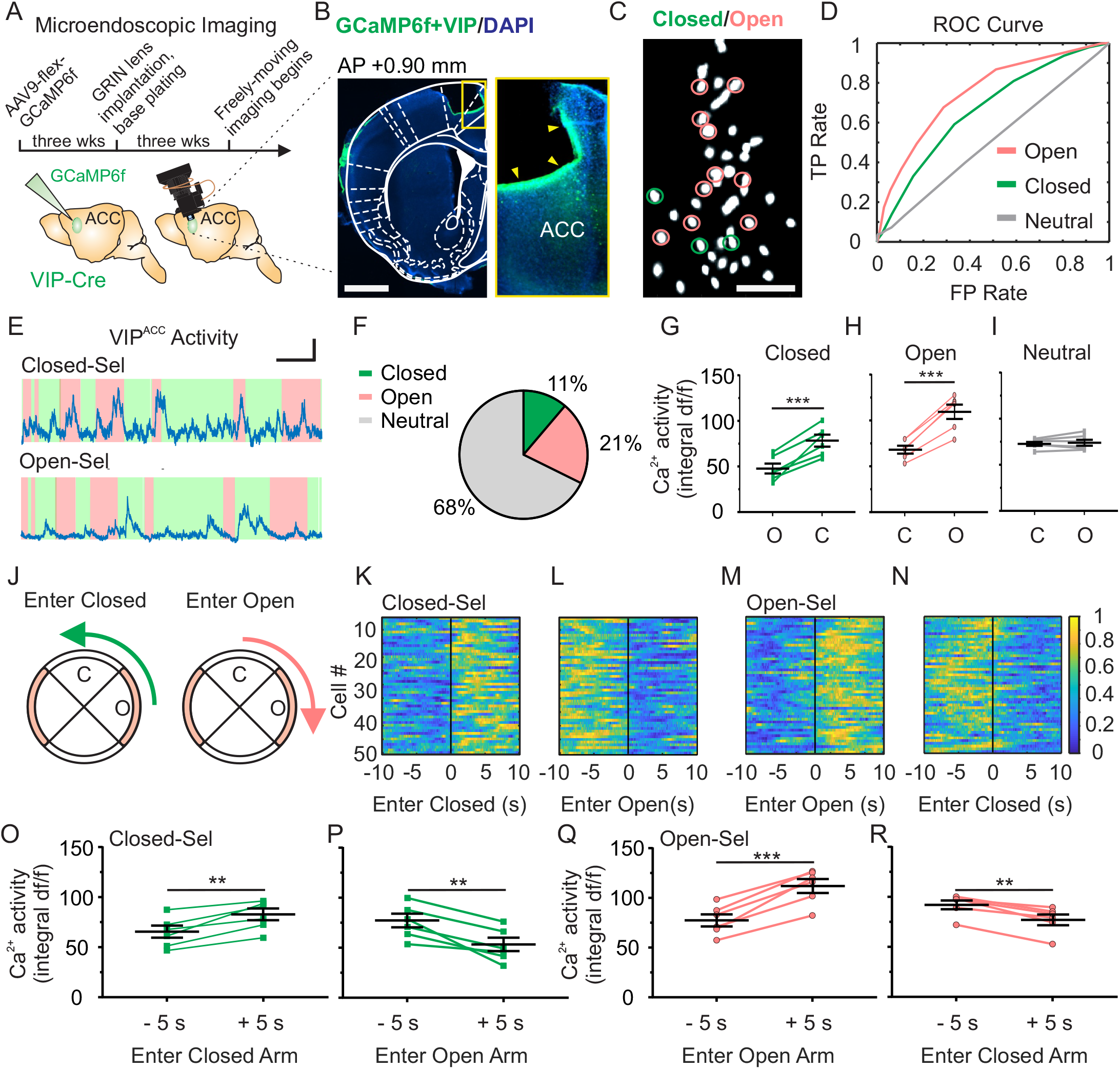
Distinct VIP^ACC^ interneurons preferentially encode anxiogenic or anxiolytic contexts in the EZM. (**A**) Experimental timeline. (**B**) Image (4x objective) showing location of the GRIN lens and GCaMP6f expression. Blue: DAPI, green: VIP^ACC^ expressing GCaMP6f. Dotted white overlay, brain regions. Yellow arrowheads, GRIN lens location in the ACC. **Left**: left hemisphere. **Right**: zoomed image of yellow boxed region. Scale bar = 1mm (left panel) or 400 μm (right panel). (**C**) VIP^ACC^ expressing GCaMP6f (white) in vivo while mouse navigates the open and closed arms of the EZM. Pink circles: open-selective, green circles: closed-selective. Neutral cells are not circled. Scale bar = 100 μm. (**D**) ROC curves demonstrating how well cells encode behavioral states for open-selective (pink, auROC = 0.75), closed-selective (green, auROC = 0.66), and neutral (gray, auROC = 0.51) VIP^ACC^. (**E**) Ca^2+^ transients: closed-selective (**top**), open-selective (**bottom**). Shaded areas indicate the location of the mouse: open (pink) or closed (green) arms. Scale bar = 25 s and 2 SD. (**F**) 21% of VIP^ACC^ were classified as open-selective, 11% as closed-selective, and 68% as neutral. (**G-I**) Ca^2+^ activity of selective VIP^ACC^ per mouse. (**G**) Closed-selective. Closed vs. open. \*\**p*=0.0016. (**H**) Open-selective. Open vs. closed. \*\*\**p*=0.0002. (**I**) Neutral cells in the EZM. Closed vs. open. *p*=0.5239. (**J**) Movement of a mouse from an open to a closed arm (**left**) or a closed to an open arm (**right**) of the EZM. (**K-L**) Activity of closed-selective cells from 10 s prior to 10 s after entering either a closed (**K**) or an open (**L**) arm. (**M-N**) Activity of open-selective from 10 s prior to 10 s after entering either an open (**M**) or closed (**N**) arm. (**O-R**) Average of selective-cell Ca^2+^ activity per mouse. (**O**) Closed-selective entering closed arm. −5 vs. +5 s. \*\**p*=0.0046. (**P**) Closed-selective entering open arm. −5 vs. +5 s. \*\**p*=0.0056. (**Q**) Open-selective entering open arm. −5 vs. +5 s. \*\*\**p*=0.0004. (**R**) Open-selective entering closed arm. −5 vs. +5 s. \*\**p*=0.0015. N = 6 mice, n = 345 cells. All statistics performed with Paired t-test. For all figures, Ca^2+^ activity refers to the area under the curve (integral df/f). All ROC curves, traces, and images are representative. Each replicate in G-I and O-R represents one mouse. C: closed arm, O: open arm, Closed-sel: closed selective, Open-sel: open-selective, df/f: Deltaf/f.

The elevated zero maze (EZM) was used to quantify anxiety-like behavior – closed arms are anxiolytic and open arms are anxiogenic (Fig 1, Fig S2). Implanted mice spent 65% less time in the open arms than in the closed, similar to unimplanted controls (Fig S2E-F and N). We monitored Ca^2+^ dynamics while animals navigated the EZM to determine whether exploration of anxiogenic areas altered VIP^ACC^ activity. We hypothesized that individual neurons would activate preferentially to either anxiolytic or anxiogenic zones. To test this, we isolated Ca^2+^ transients from each VIP^ACC^ and performed ROC analysis ((41)Fig 1D). auROC values quantify how well each cell’s activity matches a behavioral state; values near 0.5 are expected for cells that do not encode behavioral states and higher auROCs reflect better encoding of this information. For each neuron, we calculated an auROC for the closed arms and an auROC for the open arms. We identified individual VIP^ACC^ that demonstrated selectively increased activity in one arm type and classified them as open-selective (21%), closed-selective (11%), or neutral (which activated equally in both zones, 68%) (Fig 1C-F, Fig S3). The activity of individual selective cells better encoded anxiolytic states, as compared to neutral cells (Fig 1D). Closed-selective cells showed increased activity while animals were in the closed, as compared to the open arms (Fig 1E, top and G), whereas open-selective cells showed the opposite effect (Fig 1E, bottom and H).

To determine whether selective VIP^ACC^ activity rapidly changes in anxiolytic or anxiogenic contexts, we isolated trials when animals transitioned from one arm to the other (Fig 1J) and plotted VIP^ACC^ activity. Heatmaps and Ca^2+^ activity showed robust differences during behavioral transitions (Fig 1K-R). Closed-selective cells preferentially activated soon after animals entered the closed arms (27% increase, Fig 1K and O) or prior to entering the open arms (45% increase, Fig 1L and P). Open-selective cells were preferentially active soon after animals transitioned into the open arms (22% increase, Fig 1M and Q) or before leaving the closed arms (44% increase, Fig 1N and R).

We verified that this analysis method was reliable by using half of our data to identify selective cells and using those classifications to perform Ca^2+^ activity analysis on the remaining half (Fig S5A). In this second half of the data, activity of selective cells increased when mice were in the preferred context, relative to the non-preferred (23%, Fig S5B). Additionally, the majority of auROC values were near 0.5 for both zones (Fig S3A-B), suggesting that most VIP^ACC^ were neutral and did not preferentially activate to a specific context (Fig 1I, Fig S3C).

We next assessed whether these effects were context-specific, or they could also be identified in a second task of anxiety-like behavior. Similar to our findings in the EZM, we found that mice in the open field (OF) avoided the anxiogenic center zone (Fig S2C and D) and we identified zone-specific cells that encoded the animal’s behavioral state (Fig S4). In contrast to the OF, in the EZM there was no relationship between the animal’s velocity and VIP^ACC^ activity (Fig S2I and J), which suggests that the activity differences in EZM reflect the animal’s anxiogenic state, rather than signals related to locomotion. These data suggest that our selectivity classifications accurately reflect the relationship between VIP^ACC^ activity and anxiety-related behaviors.

Despite differences in behavior and the activity of individual neurons, there was no difference in the average activity of all VIP^ACC^ as mice explored different zones of the EZM or OF (Fig S2B and G-H), which suggests that VIP^ACC^ do not uniformly activate in anxiogenic or anxiolytic contexts. Additionally, the behavior of implanted, miniscope-mounted animals was compared to control mice and no behavioral differences were found, suggesting that neither surgeries nor implants induced changes in locomotion or anxiety-like behaviors (Fig S2K-N). Overall, these results support our hypothesis that VIP^ACC^ are a heterogeneous population, where subpopulations preferentially activate in either anxiolytic or anxiogenic contexts.

### Distinct subpopulations of VIP^ACC^ encode social and non-social stimuli

We next investigated whether individual VIP^ACC^ were selective for social or non-social stimuli by recording their activity as mice interacted with these stimuli over three days (Fig 2A and D). In Sociability, mice explored a chamber with an empty mesh cup (which they had been acclimated to) and a littermate in a mesh cup; in Social Novelty, the cups housed either a littermate or a novel mouse. Implanted mice spent more time with their littermate than an empty cup (Fig 2B, 2.4 fold-change), and more time with the novel mouse than the littermate (Fig 2C, 1.8 fold change), suggesting an interest in socialization and a preference for novel social stimuli.

**Figure 2.**
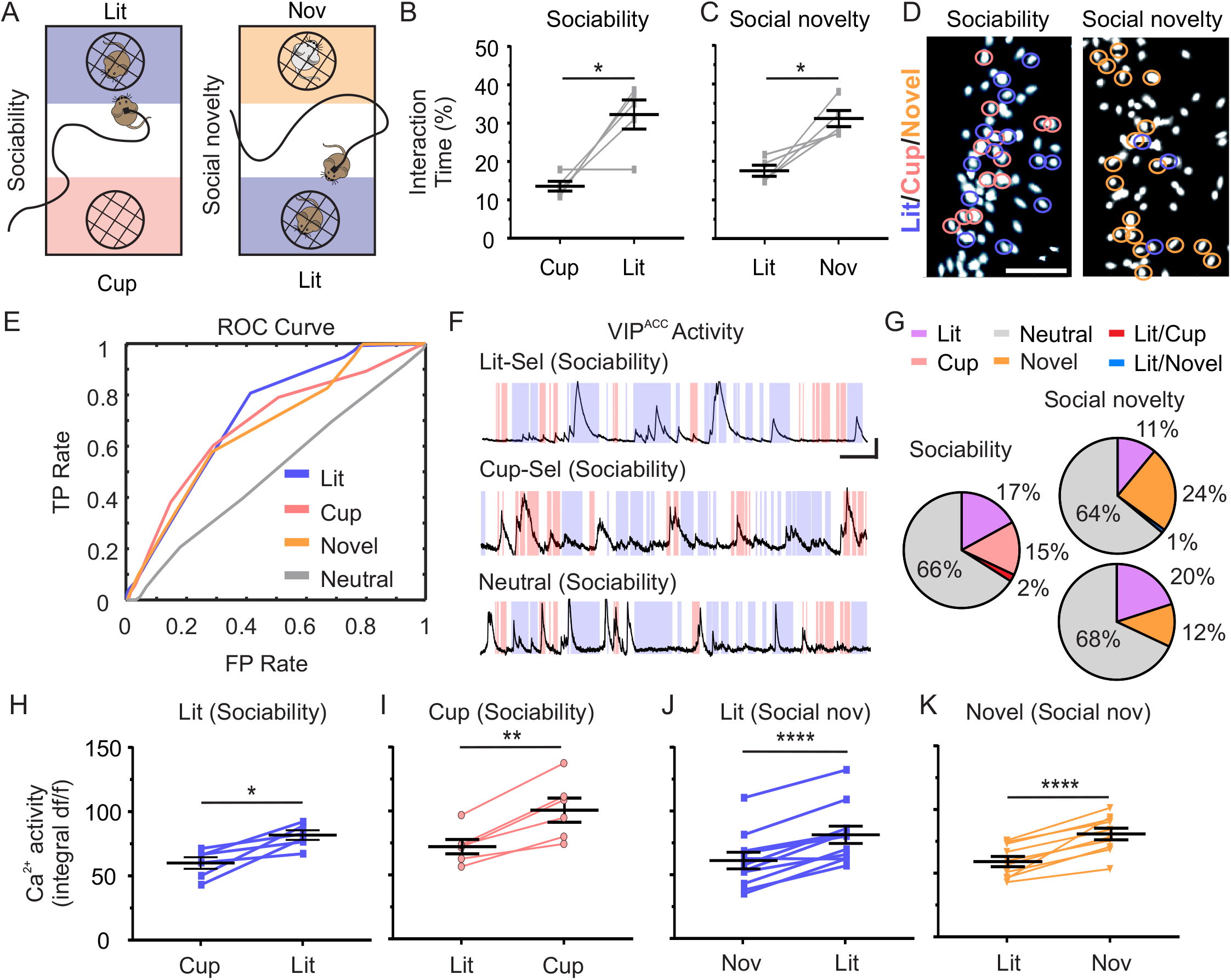
Distinct subpopulations of VIP^ACC^ encode social and non-social stimuli. (**A**) Behavioral paradigm. **Left**: Sociability (Day 1). **Right**: Social Novelty (Days 2/3). Littermate zone (purple). Empty cup zone (pink). Novel mouse zone (orange). Neutral zone (white). (**B**) Sociability. Interaction Time (%). Cup vs. littermate. \**p*=0.0199. (**C**) Social Novelty. Interaction Time (%). Littermate vs. novel mouse. \**p*=0.0141. (**D**) Images of VIP^ACC^ expressing GCaMP6f (white) in vivo during Sociability (**left**) or Social Novelty (**right**). Pink circles: cup-selective, purple circles: littermate-selective, orange circles: novel mouse-selective. Neutral cells are not circled. Scale bar = 100 μm. (**E**) ROC curves for littermate-selective (purple, auROC = 0.72), novel-mouse-selective (orange, auROC = 0.67), cup-selective (pink, auROC = 0.69), and neutral (gray, auROC = 0.51) VIP^ACC^. (**F**) Ca^2+^ transients: littermate-selective (**top**), cup-selective (**middle**), neutral cell (**bottom**). Shaded areas indicate location of mouse: littermate (purple), cup (pink), neutral (white) zones. Scale bars = 25 s and 1 SD. (**G**) In Sociability, 17% of VIP^ACC^ were classified as cup-selective, 15% as littermate-selective, 2% as selective for both cup and littermate, and 66% as neutral. In Social Novelty, VIP^ACC^ were classified as littermate-selective (11% Day 2, 20% Day 3), novel-mouse-selective (24% Day 2, 12% Day 3), selective for both littermate and novel (1% Day 2, 0% Day 3), and neutral (64% Day 2, 68% Day 3). (**H-K**) Average Ca^2+^ activity of selective cells per mouse. (**H**) Littermate-selective. Sociability. Littermate vs. cup. \**p*=0.0132. (**I**) Empty cup-selective. Sociability. Cup vs. littermate. \*\**p*=0.0017. (**J**) Littermate-selective. Social Novelty. Littermate vs. novel mouse. \*\*\*\**p*<0.0001. (**K**) Novel-mouse-selective. Social Novelty. Littermate vs. novel mouse. \*\*\*\**p*<0.0001. N = 6 mice, n = 310 cells for Sociability, n = 350 cells for Social Novelty Day 2, and n = 232 cells for Day 3. All ROC curves, traces, and images are representative. Each replicate in B-C and H-K represents one mouse. All statistics performed with Paired t-test. Lit: littermate, Nov: novel mouse, Lit-sel: littermate-selective cell, Cup-sel: cup-selective cell. Social nov: Social Novelty, df/f: Deltaf/f.

**Figure 3.**
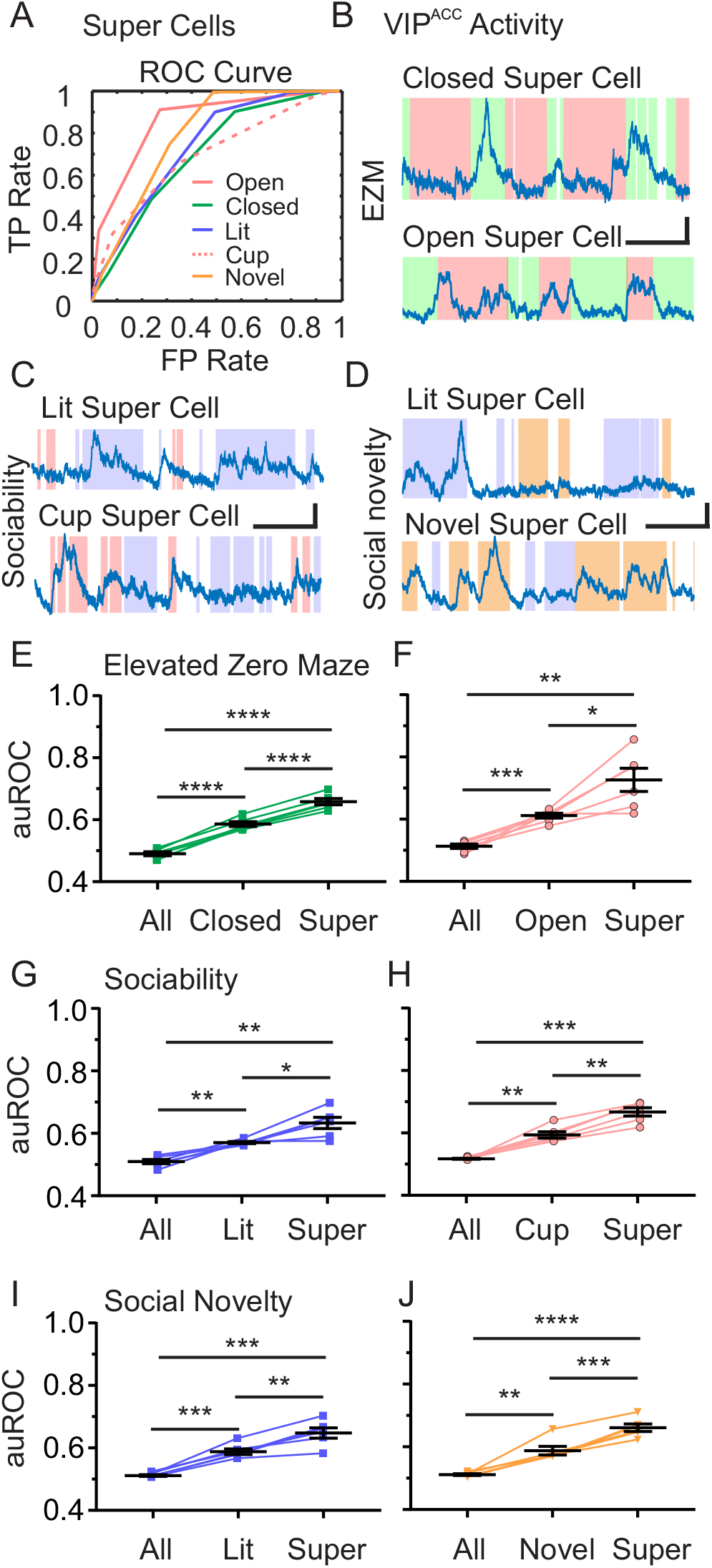
Population coding of selective VIP^ACC^ improves behavioral encoding. (**A**) ROC curves of open- (pink solid, auROC = 0.86), closed- (green, auROC = 0.70), littermate- (purple, auROC = 0.74), empty cup- (pink dotted, auROC = 0.70), and novel-mouse-selective (orange, auROC = 0.79) super cells. (**B-D**) Ca^2+^ transients of each type of super cell for the EZM (**B**) and social interaction tasks (**C-D**). Scale bars = 10 s and 1 SD. Shaded areas represent location of the mouse. Closed arm (**B**, green), open arm (**B**, pink), littermate (**C-D**, purple), empty cup (**C**, pink), novel mouse (**D**, orange) or neutral (**C-D**, white) zones. Ca^2+^ signals overlayed. (**E-J**) auROC of all cells (All), selective cells, and super cells (Super) per mouse in the EZM (**E-F**) and social interaction tasks (**G-J**). In each task, selective cells had higher auROC than the whole population and super cells had higher auROC values than selective cells. (**E-F**) This pattern was consistent in the EZM for both closed- (**E**, All vs. closed: \*\*\*\**p*<0.0001. All vs. super: \*\*\*\**p*<0.0001. Closed vs. super: \*\*\*\**p*<0.0001) and open-selective cells (**F**, All vs. open: \*\*\**p*=0.0009. All vs. super: \*\**p*=0.0047. Open vs. super: \**p*=0.0370). (**G-H**) In Sociability, this pattern was consistent for littermate- (**G,**All vs. littermate: \*\**p*=0.0022. All vs. super: \*\**p*=0.0011. Littermate vs. super: \**p*=0.0326) and cup-selective (**H,**All vs. cup: \*\**p*=0.0026. All vs. super: \*\*\**p*=0.0003. Cup vs. super: \*\**p*=0.0010). (**I-J**) In Social Novelty, this pattern was consistent for littermate- (**I**, All vs. littermate: \*\*\**p*=0.0005. All vs. super: \*\*\**p*=0.0008. Littermate vs. super: \*\**p*=0.0073.) and novel-mouse-selective (**J**, All vs. novel: \*\**p*=0.0059. All vs. super: \*\*\*\**p*-0.0001. Novel vs. super: \*\*\**p*=0.0007). \*\*\*\**p*-0.0001. N = 6 mice, n = 345 cells for EZM, n = 310 cells for Sociability, n = 350 cells for Social Novelty Day 2, n = 232 cells and Day 3. All ROC curves and traces are representative. Each replicate in E-J represents one mouse. All statistics performed with Repeated measures ANOVA with Tukey’s post test. Lit: littermate, Soc: social.

Using ROC analysis, we identified cup-, littermate-, and novel-mouse-selective cells and demonstrated that they better encode behavior states than neutral cells (Fig 2D-F, Fig S6E-F). For Sociability, cells were classified as cup-selective (15%), littermate-selective (17%), selective for both stimuli (2%), or neutral (Fig 2D-G). For Social Novelty, cells were classified as littermate-selective (11% Day 2, 20% Day 3), novel mouse-selective (24%, 12%), or neutral (64%, 68%, Fig 2D-G). Very few cells were selective for both stimuli (Fig 2G, 0-1% of cells, Littermate- and cup-selective or littermate- and novel mouse-selective cells). Littermate-selective cells showed increased activity while animals interacted with the littermate, as compared to the cup (36% increase, Fig 2F, top and H) or novel mouse (33% increase, Fig 2J). Cup-selective VIP^ACC^ were more active when animals interacted with the cup than with the littermate (40% increase, Fig 2F, middle and I). Novel mouse-selective neurons demonstrated increased activity while mice interacted with novel mice than with littermates (36% increase, Fig 2K). Similar to the OF and EZM (Fig 1I, Fig S3 and Fig S4J-M), most cells were classified as neutral and their activity did not change as mice interacted with various stimuli (Fig 2F, bottom and Fig S6A-F). We also verified that this analysis method was reliable as described above (Fig S5A) and the Ca^2+^ activity of selective cells increased when mice interacted with preferred stimuli (preferred vs. non-preferred, 17%) (Fig S5C). Similar to the anxiety-related assays, we found no differences in the average activity of all VIP^ACC^ as mice interacted with the different social or non-social stimuli (Fig S6G-H). This suggests that there is no global change in the VIP^ACC^ activity during interactions with other mice or objects. These data also suggest that, in addition to encoding for anxiogenic and anxiolytic contexts, distinct subgroups of VIP^ACC^ encode for interactions with objects, other mice, and social novelty.

### Population coding of selective VIP^ACC^ improves behavioral encoding

Our analysis thus far demonstrated that individual VIP^ACC^ can reliably encode diverse stimuli. To determine whether subsets of selective VIP^ACC^ encode behavioral states as a population to increase the reliability of their information, we averaged the Ca^2+^ traces from all cells that were selective for a specific stimulus to create “super cells” (Fig 3A-D). This process determines whether selective cells are co-activating or whether they primarily activate asynchronously. Unsynchronized Ca^2+^ transients in averaged traces would cancel each other out, resulting in auROC values similar to individual selective cells. If cells co-activate, however, the auROC values of averaged traces would be significantly higher than individual auROC values. We compared super cell auROC values to those from all VIP^ACC^ (Fig 3A, obtained by averaging the auROC values for all cells) and from the selective cells (obtained by averaging the individual auROC values of selective cells). In the EZM, the auROC values for closed super cells increased by 34% and 12% relative to all or closed-selective VIP^ACC^, respectively (Fig 3B, E). Similarly, for open super cells, we found increases of 42% and 19% in auROC values relative to all or open-selective VIP^ACC^, respectively (Fig 3B, F). We found this pattern in other behavioral tasks, as well. In Sociability, littermate super cells exhibited a 24% and 11% increase in auROC values relative to all or littermate-selective VIP^ACC^, respectively (Fig 3C, G). In Social Novelty, auROC values increased by 27% and 10% relative to all or littermate-selective VIP^ACC^, respectively (Fig 3D, I). Cup super cell auROCs increased by 29% and 13% relative to all or cup-selective VIP^ACC^, respectively (Fig 3C, H), and novel mouse super cells exhibited 24% and 11% increases in auROC values relative to all or novel-mouse-selective VIP^ACC^, respectively (Fig 3D, J). Overall, these data support previous findings and demonstrate that averaging the activity of stimulus-selective VIP^ACC^ increases the accuracy of their coding, suggesting that these cells encode information as disinhibitory subclusters.

### VIP^ACC^ subpopulations are non-overlapping and are recruited during distinct behaviors

To determine whether diverse neuronal representations are embedded in particular VIP^ACC^ subpopulations, we monitored the same cells across tasks (Fig 4A and B). By registering our neural data, we identified the same 127 neurons across tasks (out of 655 total neurons) and tracked their activity in both EZM and Sociability (Fig 4A and B). We identified distinct subpopulations of VIP^ACC^ that showed stimulus selectivity in only one task and others that were selectively active in specific zones of each task (Fig 4C). 27% of registered VIP^ACC^ were selective in the EZM and neutral during Sociability (Fig 4C). Similarly, about 22% were selective in Sociability, but neutral in the EZM (Fig 4C). Out of all registered VIP^ACC^, 17% were only open-selective, 9% only closed-selective, 16% only littermate-selective, and 6% only cup-selective. Taken together, these data show that about half of all registered VIP^ACC^ were only selective for one of these 4 stimuli, whereas only one sixth were selective for 2 or 3 stimuli (e.g., both when the mouse was in the open arm of the EZM and when it interacted with the littermate in Sociability) (Fig 4C). This demonstrates that, on average, more cells were selective for one stimulus than for two or three (Fig 4C-D). Unlike in the previous data without registration, where the majority of cells were neutral, only 36% of cells were neutral in both tasks (Fig 4C). These data suggest that, within the ACC, there are non-overlapping VIP interneuron subcircuits dedicated to processing particular stimuli.

**Figure 4.**
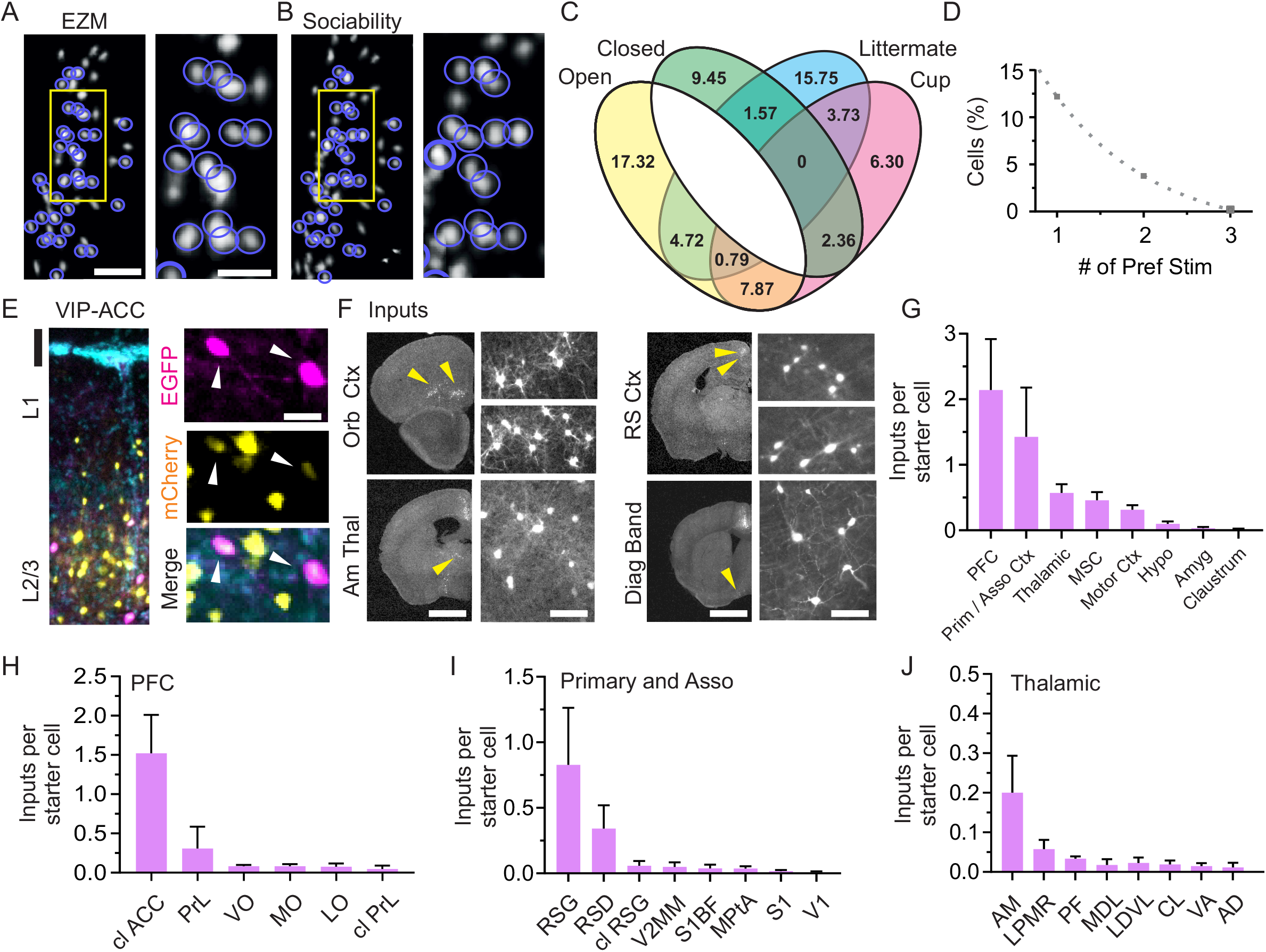
VIP^ACC^ subpopulations are non-overlapping and are recruited during distinct behaviors. (**A-B**) VIP^ACC^ were registered across tasks. Purple circles: registered cells monitored during both EZM (**A**) and Sociability **(B**). (**A-B**) **Left**: Scale bar = 100 μm. **Right**: zoomed versions of yellow outlined regions in left panels. Scale bar = 40 μm. (**C**) Venn diagram demonstrating cell selectivity in the EZM and Sociability. Each number indicates what percentage of the registered cells belong to each selectivity category. There is a gap between open and closed because these classifications are mutually exclusive – by definition a cell cannot be selective for both. (**D**) Percentage of cells selective for one, two, or three stimuli. Dotted line, one phase decay, half-life of 0.81. (**E**) Image of starter VIP^ACC^ expressing both EGFP (pseudo colored magenta, also expressing TVA and ΔG) and mCherry (pseudo colored yellow, infected by RV*dG)*, and retrogradely-labeled input cells only expressing mCherry. Coronal section. **Left**: overlay of both fluorophores and DAPI (cyan). Scale bar = 50 μm. **Right**: zoomed in superficial layers showing overlap (starter cells, white arrows). Scale bar = 15 μm. (**F**) Images of retrogradely-labeled input neurons (white) in the orbitofrontal (Orb) cortex (Ctx), anterior thalamic nuclei (Am Thal or AM), retrosplenial (RS) cortex, and diagonal band of Broca (Diag Band, part of the medial septal complex, MSC). **Left**panels: left hemisphere. Scale bar = 1.5 mm. **Right** panels: zoomed views of the retrogradely-labeled brain regions (yellow arrowheads in left panel). Scale bar = 100 μm. (**G-J**) Regions that are the most highly connected to VIP^ACC^. Rabies trans-synaptic mapping revealed that the prefrontal Ctx (PFC), primary and association Ctx (Prim/asso), thalamic regions, and MSC are highly connected to VIP^ACC^. (**H-J**) Highly connected areas (PFC, **H**, Prim/asso, **I**, thalamic, **J**) divided into subregions. Error bars represent SEM. N = 6 mice for Ca^2+^ imaging, n = 345 cells for EZM, n = 310 cells for Sociability. For tracing experiments, N = 3 mice with n = 691 starter cells and 10107 input cells. All images are representative. Pref Stim: preferred stimuli, Hypo: hypothalamus, Amyg: amygdala, cl: contralateral, PrL: prelimbic Ctx, VO: ventral orbital Ctx, MO: medial orbital Ctx, LO: lateral orbital Ctx, RSG: restrosplenial granular Ctx, RSD: retrosplenial dysgranular Ctx, V2MM: secondary visual Ctx mediomedial area, S1BF: primary somatosensory Ctx barrel field, MPtA: medial parietal association Ctx, S1: primary somatosensory Ctx, V1: primary visual Ctx, LPMR: lateral posterior thalamic nucleus medio rostral part, PF: parafascicular thalamic nucleus, MDL: mediodorsal thalamic nucleus lateral part, LDVL: laterodorsal thalamic nucleus ventrolateral part, CL: centrolateral thalamic nucleus, VA: ventral anterior thalamic nucleus, AD: anterodorsal thalamic nucleus.

### Whole-brain mapping of inputs to VIP^ACC^

To determine whether VIP^ACC^ receive projections from other brain areas involved in anxiety and social behavior, we used rabies virus-mediated trans-synaptic mapping. This technique allowed us to retrogradely label only neurons that synapse onto VIP^ACC^ (Fig 4E-J). First, we injected AAV-TVA-Glyco-GFP, which expresses target proteins in a Cre-dependent manner, in VIP-Cre mice to target VIP^ACC^. Four weeks later, we injected RV*dG*, such that starter neurons expressed both EGFP (pseudo colored magenta, also expressing TVA and ΔG) and mCherry (pseudo colored yellow, infected by RV*dG)* and input cells only expressed mCherry (Fig 4E and F). We identified these labeled cells, quantified their numbers in each brain region (Fig 4E and F), and normalized input cells to starters. We found retrogradely-labeled neurons were most prominent in other regions of the PFC (about 18% of all labeled regions), primary and associative areas (prim/asso), thalamic nuclei, and the medial septal complex (MSC), suggesting that these regions were highly connected to VIP^ACC^ (Fig 4F). We further partitioned the regions with the greatest number of retrogradely-labeled neurons into subregions and determined that VIP^ACC^ received connections from the contralateral (cl) ACC (77% of retrogradely-labeled PFC neurons, Fig 4H), prelimbic cortex (PrL, 12% of retrogradely-labeled PFC neurons, Fig 4H), retrosplenial cortex (RS, including RS granular (RSG) and dysgranular (RSD), 86% of retrogradely-labeled prim/asso neurons, Fig 4I), anteromedial thalamic nucleus (AM) (46% of retrogradely-labeled thalamic neurons, Fig 4J), and lateral posterior thalamic nucleus medio rostral part (LPMR, 8% of retrogradely-labeled thalamic neurons, Fig 4J). Our data showed that VIP^ACC^ receive long-range projections, not only from regions implicated in emotional regulation and social cognitive behavior, but also areas important for neuromodulation, memory formation, and motor actions.

## DISCUSSION

### Multimodal disinhibitory circuits in the ACC

Like many prefrontal cortical regions, the ACC is implicated in a wide range of behavioral functions. Our work supports past findings by connecting ACC activity with anxiety-related and social behavior and identifies a neural substrate for processing of multimodal stimuli. However, an important facet of our finding is that a diverse range of neuronal representations are encoded in subgroups of VIP^ACC^, which points to the varied roles of the ACC.

Human imaging studies have shown that ACC activity increases as individuals perform social and emotional tasks, as compared to non-social tasks (44). Interestingly, this previous work demonstrates heterogeneous activation of different subregions within the ACC, where increased activity of the perigenual ACC and rostral ACC is implicated in social tasks, whereas dorsal ACC is selectively activated in non-social cognitive tasks (44). Our histological analysis revealed that lenses were located at the border of the dorsal and ventral subregions of the ACC (A24a and A24b subregions). Future research could monitor VIP cells in different ACC subregions to determine whether the selectivity distributions differ. Additionally, we found subgroups of VIP^ACC^ that encoded interactions with objects, which supports past findings that characterized neural correlates in the ACC for object and location recognition and memory consolidation of object/place associations (14, 15). Our work shows that approximately 70% of VIP^ACC^ were neutral in the anxiogenic and social behavioral tasks. Perhaps this high prevalence of neutral cells reflects the multimodal nature of the ACC, where these cells encode other kinds of stimuli. For example, previous work highlighted the role of the ACC in pain processing (45). Alternatively, cells that were classified as neutral may be equally activated by multiple stimuli. For example, neutral EZM cells may preferentially activate to both anxiogenic and anxiolytic stimuli and be involved in encoding both.

When we averaged the activity of stimulus-selective VIP^ACC^, the reliability of their coding increased across diverse stimuli. Electrical coupling has been observed within inhibitory networks, including VIP interneurons (22, 46). VIP cells can disinhibit members of their own population, leading to increased co-activation of the population, which may allow these subnetworks to encode stimuli as a population and amplify the population output (22).

### Whole brain mapping of long-range direct inputs to VIP^ACC^

Using rabies trans-synaptic mapping, we provided a brain-wide map of inputs to VIP^ACC^. Additionally, we demonstrated that VIP^ACC^ receive extensive connections from the AM and RS, which are both known to project to the ACC (47–50). AM lesions are associated with memory impairments (51–53), so AM-ACC projections may be important for social memory. In support of this, mice lacking *Shank3* in the ACC show abnormal behavior in a social novelty task (8). Additionally, memory formation leads to induction of immediate-early genes in the ACC (54). In the auditory cortex, VIP cells facilitate learning about unexpected, salient events (55), suggesting that inhibitory circuits in the ACC could mediate specific aspects of memory formation. Although it remains to be tested, AM-VIP^ACC^ projections are positioned to mediate social learning and memory. In rodents, AM-ACC projections regulate histaminergic itch-induced scratching behavior (56), suggesting that, similar to sensory thalamocortical projections, this higher-order network could be composed of thalamic pathways for parallel processing (31). The RS plays a role in learning, memory, and navigation and is highly connected to the anterior thalamic nuclei and hippocampus (51, 57–59). Spatial information conveyed to VIP^ACC^ from the RS could guide motor actions during social behavior (58, 60). We also showed that VIP^ACC^ are highly connected to other prefrontal cortex subregions and the contralateral ACC. Our results support human imaging studies demonstrating that distinct ACC-PFC networks are involved in diverse aspects of emotional processing (1) and monitor behavior to guide compensatory systems (61).

Our injections and lenses were located at the border between the A24a and A24b subregions of the ACC (62, 63). Past research shows that A24 receives projections from the amygdala, orbitofrontal cortex, thalamus, RS, and motor cortex and moderate projections from hippocampus, hypothalamus, and autonomic brain nuclei (63–65). Although our data is mostly consistent with these findings, a major difference is that we used trans-synaptic viral mapping in a cell-type specific manner, while other studies used classic neuronal tracers to map connectivity to all ACC neurons (63–65). Mapping experiments that include inputs to other interneuron subtypes will provide a better framework to understand ACC function and the connectivity patterns of inhibitory circuits of the ACC.

### Heterogeneous inhibitory subcircuits in the ACC

VIP interneurons express a variety of neuromodulator receptors, which may allow long-range projections to exert context-dependent disinhibition (19, 20, 55, 66–69). Our data showed that subgroups of VIP^ACC^ are engaged by particular stimuli. Through the actions of neuromodulators, VIP interneurons in the ACC may recruit subpopulations of Pyr that are behaviorally relevant or encode specific information.

We demonstrated that a subgroup of VIP^ACC^ are engaged by social novelty. Locus coeruleus (LC) neurons, the primary source of norepinephrine in the forebrain, are recruited by novel stimuli, so this neuromodulator may be important in social novelty (70–73). Additionally, mice exhibit elevated arousal when exposed to novel or social stimuli, which is abolished upon lesioning of the ACC or LC (74). This suggests that noradrenergic modulation of the ACC may be a substrate for attention and novelty (74). In the cortex, norepinephrine differentially regulates the activity of interneurons (75, 76), but it is unknown whether recruitment of VIP^ACC^ by novel social stimuli is dependent on particular neuromodulators. We show that a small number of VIP^ACC^ are selective to multiple stimuli or contexts. It is possible that through the actions of neuromodulators, distinct groups of VIP^ACC^ could overlap in their activity, allowing for the co-activation of segregated Pyr subgroups (77). This cellular mechanism could enable ACC networks to bind multiple streams of information to guide behavioral actions or monitor optimal performance (60, 61, 78). In the barrel cortex, the majority of VIP interneurons are located in layer 2/3, but those in other cortical layers exhibit different dendritic and axonal morphology (79). Due to the large field of view in our Ca^2+^ imaging experiments and variability in lens placement, it was not possible to correlate the activity of VIP to different cortical layers.

Using in vivo imaging with cellular resolution in freely moving mice, we showed that VIP^ACC^ are functionally heterogeneous, where distinct subcircuits encode for diverse stimuli. ROC analysis identified stimulus-selective VIP^ACC^ that encoded anxiety-like behaviors or interactions with mice and objects, even though there was no difference in overall VIP^ACC^ activity. Averaging the activity of selective VIP^ACC^ enhanced their ability to encode for specific stimuli. We also determined that most VIP^ACC^ were either selective for only one stimulus or were neutral. Lastly, we used trans-synaptic mapping to provide a map of VIP^ACC^ connectivity and show that VIP^ACC^ receive inputs from regions implicated in emotional regulation, social cognition, and memory formation. To our knowledge, these data provide the first evidence of functional heterogeneity of VIP^ACC^ in vivo and show that population coding of selective VIP^ACC^ may encode stimulus-specific information, which provides a framework for how the ACC encodes information across diverse behavioral states.

## ACKNOWLEDGEMENTS

We thank all research assistants in the Cruz-Martín lab, as well as Margaret Minnig, Timothy Otchy, Nathan Perkins, and Daniel P. Leman for optimization of miniscope imaging and lens implant placement, Tim Gardner for providing access to 3D printers and sharing video acquisition software, Peyman Golshani and Daniel Aharoni for donating the camera sensor and data acquisition board for miniscope experiments. Ashley Comer, Nancy Padilla, Mark Howe, William A. Liberti 3rd and members of the Cruz-Martín lab helped critically by reading the manuscript and engaging in helpful discussions. Todd Blute and the Boston University Biology Imaging Core provided use of the epifluorescence microscope. Lastly, we used the Boston University Shared Computing Cluster to analyze our data. This work was supported by a NARSAD Young Investigator Grant (AC-M, #27202), the NSF NRT UtB: Neurophotonics National Research Fellowship (LNK, #DGE-1633516), and the Boston University Undergraduate Research Opportunities Program (TPHN, WWY) (https://www.bbrfoundation.org/grants-prizes/grants, https://www.nsf.gov/; http://www.bu.edu/urop/). The funders had no role in study design, data collection and analysis, decision to publish, or preparation of the manuscript.

## CONTRIBUTIONS

**Connor Johnson:** Conceptualization: formulated ideas for data analysis (equal), Software: code writing and analysis for behavioral and calcium imaging data, calcium activity analysis with CaImAn (lead). Formal Analysis: analysis of calcium data (equal). Investigation: surgeries, histology, episcope imaging, cell counting (equal). Validation: validation of calcium analysis strategy (lead). Writing – Original Draft: part of methods (support). Writing – Review & Editing: provided edits (equal). Visualization: figure generation (support).

**Lisa N. Kretsge:** Conceptualization: formulated composition and goals of the paper (equal), Software: behavioral analysis with DeepLabCut (equal). Formal Analysis: histological analysis, analysis of trans-synaptic tracing data, statistical analyses (equal). Investigation: behavioral experiments, histology, episcope imaging (equal), Writing – Original Draft: wrote initial draft (excluding part of methods and discussion) (lead). Writing – Review & Editing: incorporated edits (lead). Visualization: figure generation (equal).

**William W. Yen:** Methodology: miniscope adaptation design and construction, developed protocols for GRIN lens implant surgeries (lead). Investigation: surgeries and baseplating, behavioral experiments with miniscopes (lead). Writing – Review & Editing: provided edits (support). Visualization: figure generation (support).

**Balaji Sriram:** Conceptualization: provided ideas for analysis (support). Investigation: set up DeepLabCut with our data (support). Supervision: oversight of some analysis projects (equal).

**Jessica C. Jimenez:** Methodology: taught our lab to perform surgeries and use miniscopes (equal). Writing – Review & Editing: provided edits (equal).

**Tushare J. Jinadasa:** Formal Analysis: quantification of trans-synaptic tracing data (equal). Investigation: surgeries, histology, episcope imaging, cell counting (equal). Writing – Review & Editing: provided edits (support). Visualization: figure generation (support).

**Alexandra O’Connor:** Software: code writing and analysis of the output of DeepLabCut data (equal).

**Ruichen Sky Liu:** Software: code writing for calcium imaging analysis (support). Writing – Original Draft: part of methods (support). Visualization: activity heatmaps figure generation (support).

**Thanh P. H. Nguyen:** Methodology: miniscope adaptation design and construction (equal).

**Eun Seon Cho:** Investigation: annotation for behavioral analysis in DeepLabCut (support).

**Erelle Fuchs:** Investigation: surgeries, histology, episcope imaging, cell counting (support).

**Eli D. Spevack:** Investigation: histology, episcope imaging, cell counting (support).

**Berta Escude Velasco:** Investigation: cell counting (support). Writing – Review & Editing: provided edits (support).

**Frances S. Hausmann:** Investigation: perfusions and histology (support).

**Alberto Cruz-Martín:** Conceptualization: formulated composition, goals, and scope of the paper and approaches for analyses (lead), Formal Analysis: statistical analyses (equal). Writing – Original Draft. Writing – Review & Editing: editing and feedback throughout (lead).

Visualization: figure design and generation (lead). Supervision: mentorship and oversight of the project (lead). Project Administration: management and coordination (lead). Funding acquisition (lead).

## AVAILABILITY OF DATA AND MATERIALS

Data Availability: All relevant data are within the paper, and underlying data are available at https://github.com/CruzMartinLab. Custom-written routines for behavioral tracking, calcium imaging analysis, and miniscope models and tools are available at https://github.com/CruzMartinLab. For further information, please contact the corresponding author.

