## Supplemental table and figures for "Distinct VIP interneurons in the cingulate cortex encode anxiogenic and social stimuli"

### SUPPLEMENTARY INFORMATION

#### Table S1

| Part Name | Supplier | URL |
|---|---|---|
| UCLA Miniscope CMOS | LabMaker GmbH, Berlin, Germany | https://www.labmaker.org/products/cmos-pcb-miniscope |
| Custom filters: 470/40 EM (4mm x 4mm) (excitation), 535/50 EM (4mm x 4mm) (emission), 495nm BS (4mm x 4.8mm) (dichroic) | Chroma Technology, Cambridge, Massachusetts | |
| LUXEON Rebel Blue LED (470nm, 70 lm @ 700mA) (#LXML-PB01-0040) | Lumileds, San Jose, California | http://www.lumileds.com/products/color-leds/luxeon-rebel-color |
| Achromat lens (5mm diameter x 15mm FL, MgF2 Coated, Achromatic Doublet Lens) (#45-206) | Edmund optics, Barrington, New Jersey | http://www.edmundoptics.com/optics/optical-lenses/achromatic-lenses/mgf2-coated-achromatic-lenses/45207/ |
| Drum lens (2.4mmx3.0mm) (#45-549) | Edmund optics, Barrington, New Jersey | http://www.edmundoptics.com/optics/optical-lenses/ball-condenser-lenses/drum-lenses/45549/ |
| Objective GRIN lens (objective 2.0mm Dia, 0.25 pitch) (#GT-IFRL-200-inf-50-NC) | GRINtech, Jena, Germany | http://www.grintech.de/ |
| Objective GRIN lens (objective 1.8mm Dia, 0.25 pitch) (#64-519) | Edmund optics, Barrington, New Jersey | http://www.edmundoptics.com/optics/optical-lenses/aspheric-lenses/gradient-index-grin-rod-lenses/64519/ |
| Implanted GRIN lens (Relay Lens, 1 mm diameter, 4 mm length, 0.44 pitch length, 0.47 NA) (#GT-IFRL-100) | GRINtech, Jena, Germany | |
| UCLA DAQ board | LabMaker GmbH, Berlin, Germany | https://www.labmaker.org/products/daq-pcb |
| 50 ohm coax silicone rubber jacketed cable (#CW2040-3650SR) | Cooner Wire, Chatsworth, California | https://www.coonerwire.com/mini-coax/ |
| SMA Coax Connector (for connectorizing the coax cable above) (#901-9867-RFX) | Digikey Electronics, Thief River Falls, Minnesota | http://www.digikey.com/product-search/en?vendor=0&keywords=901-9867-RFX |
| UCLA Miniscope Acquisition Software | University of Cailfornia at Los Angeles, Los Angeles, California | http://miniscope.org/index.php?title=Main_Page |

**Table S1. Miniscope parts.** Table lists names of products, suppliers, and websites with detailed information for each purchased miniscope part. Table does not include parts that were 3D printed in-house (Fig S1).

**Figure S1**

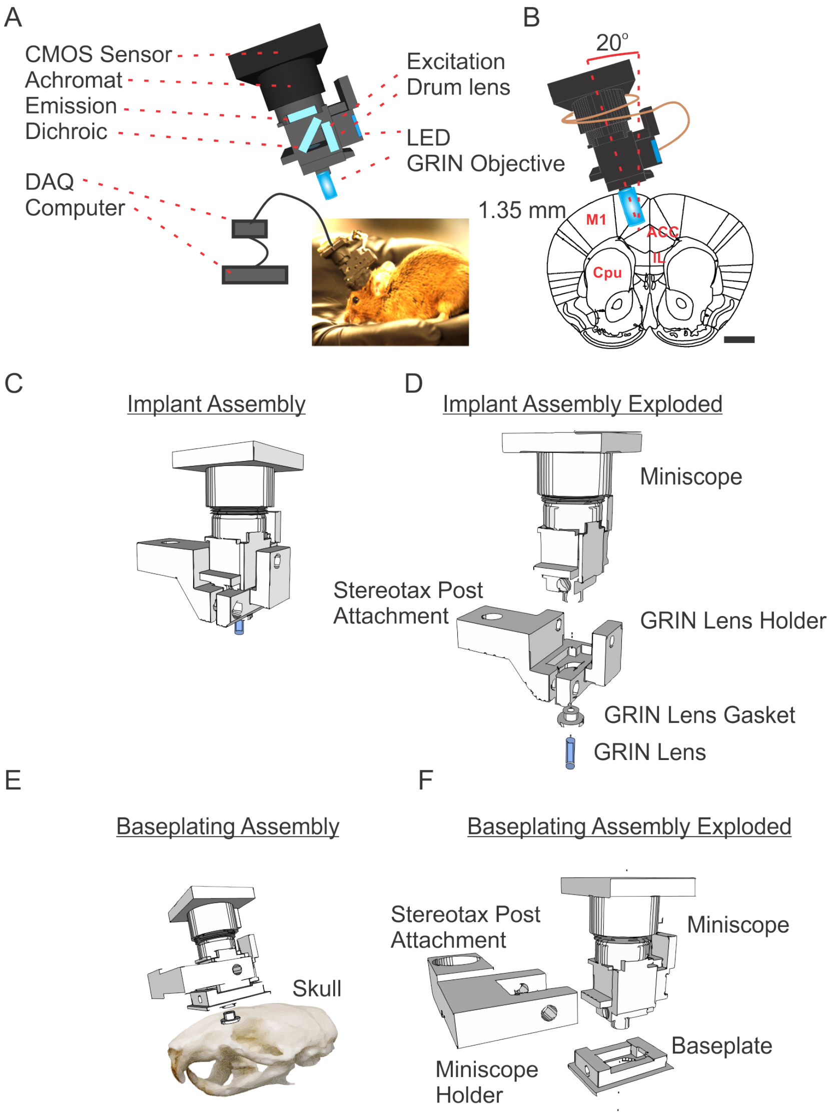


**Figure S1. 3D-printed miniscope for imaging in the ACC.** (**A**) Model displaying the main components of the miniscope and image of a miniscope implanted in an experimental mouse. (**B**) Schematic of a miniscope implanted in the ACC (coronal section, implanted at a 20 degree angle). Scale bar = 1.5 mm. (**C-D**) Implant assembly (**C**) and exploded version (**D**) showing stereotax post attachment, GRIN lens holder, and GRIN lens gasket. This is used to surgically implant a GRIN lens. (**E-F**) Baseplating assembly (**E**) and exploded version (**F**) showing stereotax post attachment, miniscope holder, and baseplate. This is used to attach baseplate. Miniscope models available at https://github.com/CruzMartinLab.

**Figure S2**

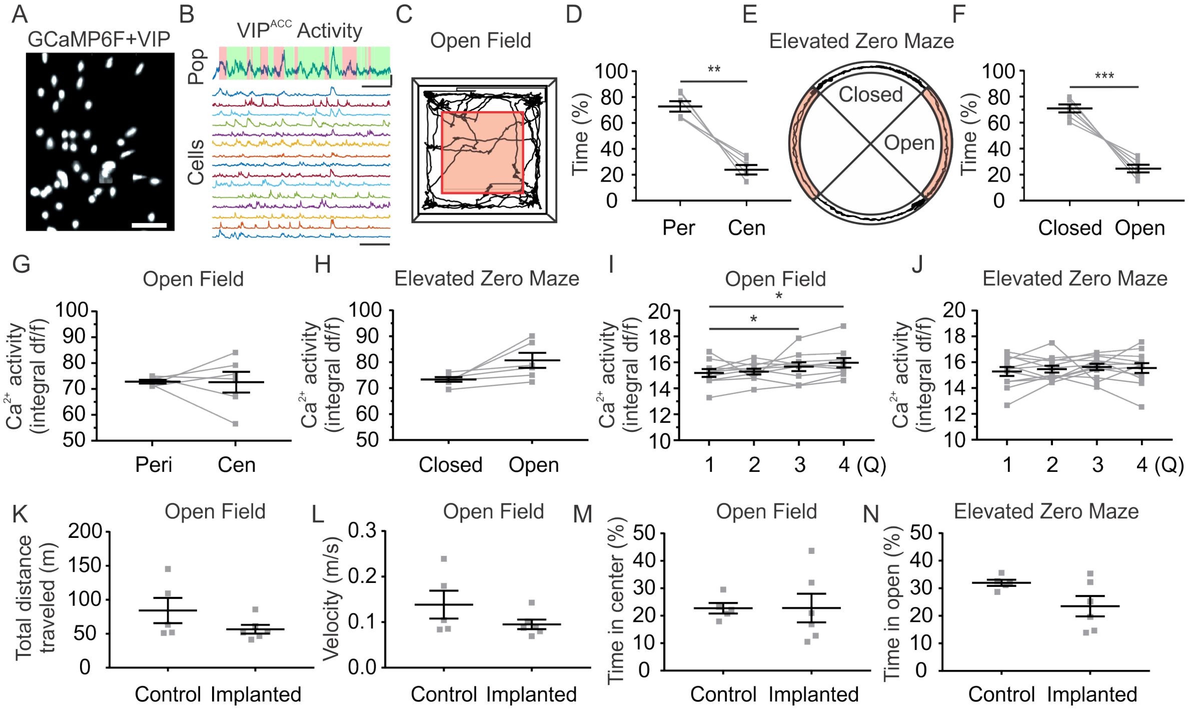


**Figure S2. Population of VIP$^{ACC}$ do not uniformly activate in anxiolytic or anxiogenic zones.** (**A**) Image of VIP$^{ACC}$ expressing GCaMP6f (white) in vivo. Scale bar = 100 µm. (**B**) **Top**: population activity of VIP$^{ACC}$ represented as SD. Location of mouse: open (pink) or closed (green) arms of the EZM. Scale bar = 50 s. Scale bar = 1 SD. **Bottom**: $Ca^{2+}$ transients (normalized to peak activity) while the animal explores the EZM. Scale bar = 50 s. (**C**) Trace of locomotor activity in the OF. Pink zone: center. Outside of the pink zone: periphery. (**D**) Time (%). OF. Periphery vs. center. ***p*=0.0014. (**E**) Trace of locomotor activity in the EZM. Pink zone: open arms. White zone: closed arms. (**F**) Time (%). EZM. Closed vs. open. ****p*=0.0006. (**G-J**) $Ca^{2+}$ activity of all VIP$^{ACC}$ per mouse. (**G**) OF. Periphery vs. center. *p*=0.9575. (**H**) EZM. Closed vs. open. *p*=0.0510. (**I**) $Ca^{2+}$ activity across 4 quartiles of velocity. OF. Repeated measures ANOVA with Tukey's test. Q1 vs Q3. **p*=0.0382. Q1 vs Q4. **p*=0.0271. **(J)** $Ca^{2+}$ activity across 4 quartiles of animal velocity. EZM. Repeated measures ANOVA with Tukey's test. All quartile comparisons. $p > 0.05$. (**K-N**) Control non-surgerized mice did not behave differently from mice with implants and miniscopes in the OF or EZM. (**K**) Distance traveled. OF. *p*=0.2209. (**L**) Velocity. OF. *p*=0.2388. (**M**) Time (%) in center. OF. *p*=0.9916. (**N**) Time (%) in open arms. EZM. *p*=0.07. N = 6 implanted mice for $Ca^{2+}$ imaging and N = 5 control mice. For OF, n = 273 cells, for EZM, n = 345 cells. All traces and images are representative. Each replicate in D and F-N represents one mouse. All statistics performed with Paired t-test unless otherwise stated. Q: quartile, df/f: Deltaf/f.

**Figure S3**

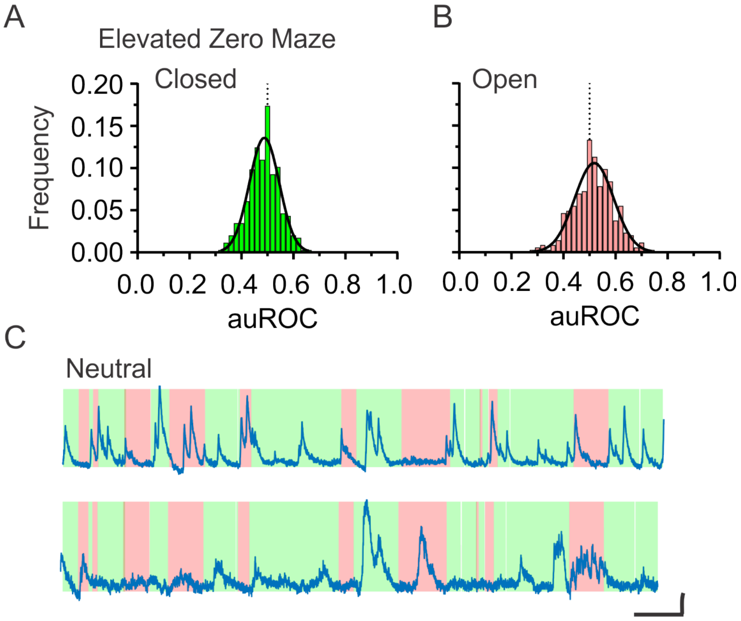


**Figure S3. Frequency distribution for auROC values in OF and EZM.** (**A-B**) Frequency distributions for auROC values all cells in the EZM for closed (**A**) or open (**B**) arms. Black curves are Gaussian fits. (**A**) Closed arms. EZM. Gaussian fit: Amplitude = 0.14, Mean = 0.49, SD = 0.06. (**B**) Open arms. EZM. Gaussian fit: Amplitude = 0.11, Mean = 0.52, SD = 0.07. (**C**) $Ca^{2+}$ traces of representative neutral cells in the EZM. Location of mouse: open (pink) or closed (green) zones. **Top**: auROC = 0.51 for closed, 0.51 for open. **Bottom**: auROC = 0.49 for closed, 0.51 for open. Scale bar = 2 SD and 25 s. N = 6 mice and n = 345 cells.

### Figure S4

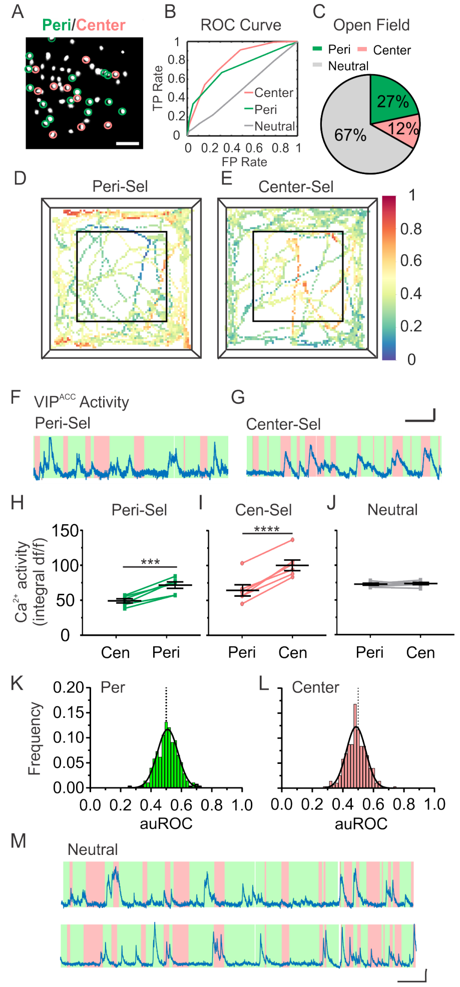


**Figure S4. Subpopulations of VIP$^{ACC}$ preferentially activate to the center and periphery of the OF.** (**A**) Image of VIP$^{ACC}$ expressing GCaMP6f (white) in vivo. Pink circles: center-selective, green circles: periphery-selective. Neutral cells are not circled. Scale bar = 100 µm. (**B**) ROC curves for center-selective (pink, auROC = 0.79), periphery-selective (green, auROC = 0.72), and neutral (gray, auROC = 0.50). (**C**) 12% of VIP$^{ACC}$ were classified as center-selective, 27% as periphery-selective, and 67% as neutral. (**D-E**) Heatmaps of VIP$^{ACC}$ activity normalized to peak activity while mice explore the OF for periphery- (**D**) and center-selective (**E**). Inner black line: border between center and periphery. Warmer colors represent greater neural activity than cooler colors. (**F-G**) $Ca^{2+}$ transients: periphery-selective (**F**) and center-selective (**G**). Shaded areas: center (pink) and periphery (green) zones. Scale bar = 25 s and 2 SD. (**H-J**) $Ca^{2+}$ activity of selective cells per mouse. (**H**) Periphery-selective cells. Center vs periphery. ****p*=0.0005. (**I**) Center-selective cells. Periphery vs. center. *****p*<0.0001. (**J**) Neutral cells. Periphery vs. center. *p*=0.8996. (**K-L**) Frequency distributions for auROC values all cells in the OF for periphery (**K**) or center (**L**). Black curves are Gaussian fits. (**K**) Periphery. OF. Gaussian fit: Amplitude = 0.12, Mean = 0.51, SD = 0.07. (**L**) Center. OF. Gaussian fit: Amplitude = 0.12, Mean = 0.49, SD = 0.06. (**M**) $Ca^{2+}$ traces of representative neutral cells in the OF. Location of mouse: center (pink) or periphery (green) zones. **Top**: auROC = 0.52 for periphery, 0.48 for center. **Bottom**: auROC = 0.47 for periphery, 0.53 for center. Scale bar = 2 SD and 25 s. N = 6 mice, n = 273 cells. All ROC curves, traces, and images are representative. Each replicate in H-J represents one mouse. All statistics performed with Paired t-test. Per: periphery, Cen: center, df/f: Deltaf/f.

**Figure S5**

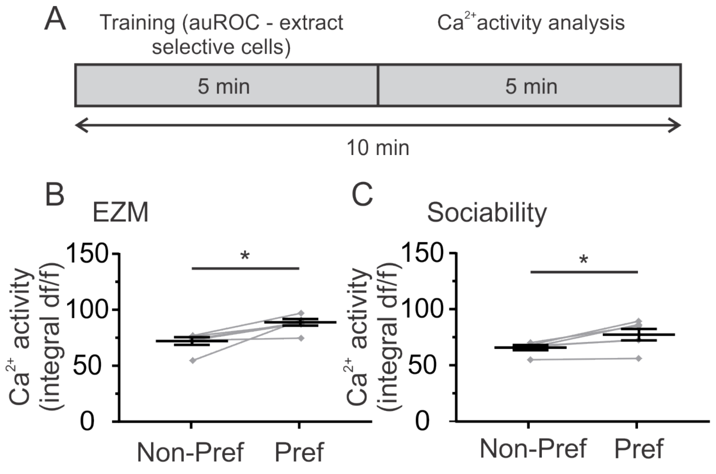


**Figure S5. Validation of VIP$^{ACC}$ activity analysis.** (**A**) Schematic explaining validation of analysis. (**B-C**) Cells identified as selective using one half of the data had increased activity in the second half of the data when mice explored the preferred stimulus. (**B**) Grouped data for EZM. Preferred vs. Non-preferred. *$p$=0.0166. N = 6 mice. (**C**) Grouped data for the social interaction task. Preferred vs. Non-preferred, *$p$=0.0216. N = 6 mice, for EZM, n = 345 cells, for Sociability, n = 310 cells. Each replicate represents the average data for one mouse. All statistics performed with Paired t-test. Non-pref: non-preferred, Pref: preferred, df/f: Deltaf/f.

**Figure S6**

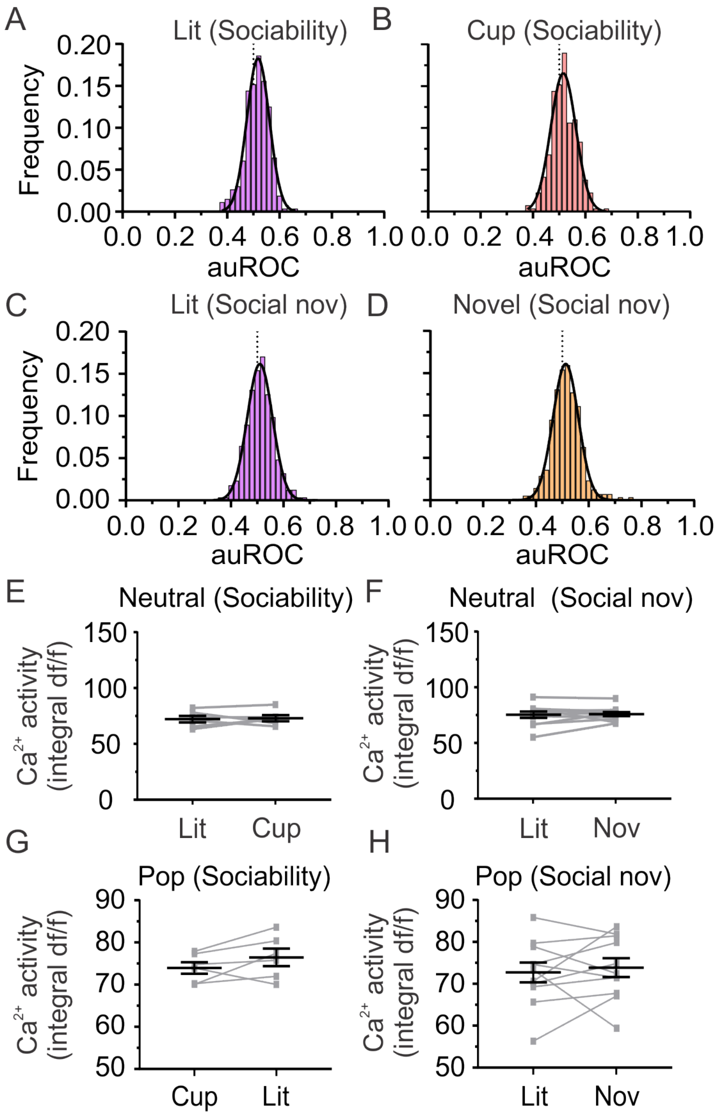


**Figure S6. Frequency distribution for auROC values in the social interaction task.** (**A-D**) Frequency distribution of auROC values in the social interaction task. Black curves: Gaussian fits. (**A**) Littermate. Sociability. Gaussian fit: Amplitude = 0.18, Mean = 0.52, SD = 0.04. (**B**) Cup. Sociability. Gaussian fit: Amplitude = 0.17, Mean = 0.52, SD = 0.05. (**C**) Littermate. Social Novelty. Gaussian fit: Amplitude = 0.16, Mean = 0.51, SD = 0.05. (**D**) Novel mouse. Social Novelty. Gaussian fit: Amplitude = 0.16, Mean = 0.51, SD = 0.05. (**E-F**) Activity of neutral cells per mouse. (**E**) Sociability. Littermate vs. cup. $p$=0.8026. (**F**) Social Novelty. Littermate vs. Novel mouse. $p$=0.7879. (**G-H**) $Ca^{2+}$ activity of population $VIP^{ACC}$ per mouse. (**G**) Sociability. Littermate vs. cup. $p$=0.1814. (**H**) Social Novelty. Novel mouse vs. littermate. $p$=0.6364. Each replicate in E-H represents one mouse. N = 6 mice, n = 310 cells for Sociability, n = 350 cells for Social Novelty Day 2, n = 232 cells and Day 3. All statistics performed with Paired t-test. Lit: littermate, Nov: novel mouse, Social nov: Social Novelty, Pop: population, df/f: Deltaf/f.
